# The lymphoid-associated interleukin 7 receptor (IL-7R) regulates tissue resident macrophage development

**DOI:** 10.1101/534859

**Authors:** Gabriel A. Leung, Taylor Cool, Clint H. Valencia, Atesh Worthington, Anna E. Beaudin, E. Camilla Forsberg

**Affiliations:** Quantitative and Systems Biology Program, University of California-Merced, Merced, CA, USA; Institute for the Biology of Stem Cells, University of California-Santa Cruz, Santa Cruz, CA, USA; Department of Biological Sciences, San Jose State University, San Jose, CA, USA; Molecular and Cell Biology Department, School of Natural Sciences, University of California-Merced, Merced, CA, USA

## Abstract

The discovery of a fetal origin for tissue-resident macrophages (trMacs) has inspired an intense search for the mechanisms underlying their development. Here, we performed in vivo lineage tracing of cells with an expression history of IL-7Rα, a marker exclusively associated with the lymphoid lineage in adult hematopoiesis. Surprisingly, we found that IL7R-Cre labeled fetal-derived, adult trMacs. Labeling was almost complete in some tissues and partial in other organs. The putative progenitors of trMacs, yolk sac (YS) erythromyeloid progenitors (EMPs), did not express IL-7R, and YS hematopoiesis was unperturbed in IL-7R-deficient mice. In contrast, tracking of IL-7Rα message levels, surface protein expression, and IL7R-Cre-mediated labeling across fetal development revealed dynamic regulation of IL-7Rα mRNA expression and rapid upregulation of IL-7Rα surface protein upon transition from monocyte to macrophage within fetal tissues. Fetal liver monocyte differentiation *in vitro* produced IL-7R+ macrophages, supporting a direct progenitor-progeny relationship. Additionally, blockade of IL-7R function during late gestation specifically impaired the establishment of fetal-derived tissue macrophages in vivo. These data provide evidence for a distinct function of IL-7Rα in fetal myelopoiesis and identify IL-7R as a novel regulator of tissue-resident macrophage development.

## Introduction

Hematopoietic stem cells (HSCs) are responsible for sustaining blood and immune cell production across the lifespan of the animal, under steady-state conditions, during infection, and following transplantation. However, recent findings have revealed that adult HSCs have a limited capability to generate or regenerate many tissue-resident immune cells, including subsets of tissue-resident macrophages (trMacs) such as microglia, epidermal Langerhans cells, liver Kupffer cells, and alveolar macrophages (Beaudin and Forsberg, 2016; Cool and Forsberg, 2019; Ginhoux et al., 2010; Gomez Perdiguero et al., 2015; Guilliams et al., 2013; Hashimoto et al., 2013; Hoeffel et al., 2012; Sawai et al., 2016; Yona et al., 2013). TrMacs are specialized macrophages that reside within tissues and have specific functions in tissue and immune homeostasis (Epelman et al., 2014; Ginhoux et al., 2010; Gomez Perdiguero et al., 2015; Guilliams et al., 2013; Hoeffel et al., 2012). Unlike classical adult monocyte-derived macrophages that circulate and have high turnover rates, trMacs are maintained via local proliferation within specific tissues, largely independent of contribution from adult hematopoiesis (Ajami et al., 2007; Hashimoto et al., 2013; Hulsmans et al., 2017; Merad et al., 2002). Recent fate mapping studies have provided direct evidence for a fetal origin of specific trMacs (Epelman et al., 2014; Gomez Perdiguero et al., 2015; Hoeffel et al., 2015; Schulz et al., 2012; Yona et al., 2013). However, the cellular and molecular mechanisms driving trMac establishment and expansion within fetal tissues are poorly understood.

Accumulating evidence points towards extraembryonic yolk sac (YS)-derived erythromyeloid progenitors (EMPs) as the initial cell-of-origin for macrophages that seed developing tissues, and the primary source of brain microglia (Ginhoux et al., 2010; Gomez Perdiguero et al., 2015; Kierdorf et al., 2013). Microglia and other trMacs can differentiate directly from EMPs through a myb-independent pathway or through a myb-dependent pathway, likely through an EMP-derived intermediate (Hoeffel et al., 2015; Schulz et al., 2012). In certain fetal-derived trMac populations, including lung alveolar macrophages, epidermal Langherhans cells, and liver Kupffer cells, macrophages derived from fetal liver (FL)-derived precursors may replace or replenish initially seeded YS-derived macrophages (Guilliams et al., 2013; Hoeffel et al., 2015; Yona et al., 2013). However, both the precise cell of origin for FL precursors and their specific contribution to adult trMac compartments remain controversial, due in part to incomplete progenitor labeling and the difficulties in accurately tracking progeny in situ. Despite intense investigation of the mechanisms regulating macrophage differentiation from YS progenitors, including gene expression programs (Mass et al., 2016) and mechanisms of tissue seeding (Stremmel et al., 2018), comparatively less is known about the developmental mechanisms that regulate the establishment of tissue macrophages from later waves of hematopoietic cell production.

Here, our investigation of hematopoietic development using the IL7R-Cre lineage tracing model (Schlenner et al., 2010) revealed unexpected and robust labeling of adult trMacs in multiple tissues. Examination of fetal myeloid development revealed a transient wave of IL-7Rα expression on developing fetal macrophages that was dynamically regulated as macrophages differentiated within developing resident tissues. Both germline deletion and specific blockade of the IL-7R during the developmental window in which fetal tissue macrophages express IL-7R confirmed the functional requirement for IL7R signaling during fetal trMac development. In contrast, IL-7Rα was not expressed by YS EMPs and yolk sac hematopoiesis was unperturbed in IL7R^−/−^ embryos. Together, these experiments reveal IL-7R as a novel regulator of fetal macrophage development during late gestation.

## Results

### IL7R-Cre specifically labels adult tissue-resident macrophages

The lymphoid-associated genes Flk2, IL-7Rα and Rag1 label cells with increasingly restricted lymphoid potential in adult hematopoiesis (Adolfsson et al., 2005; Forsberg et al., 2006; Igarashi et al., 2002; Kondo et al., 1997). In contrast, all three mark oligopotent hematopoietic progenitors with both myeloid and lymphoid potential during fetal development (Beaudin et al., 2016; Boiers et al., 2013). While we and others have used the Flk2-Cre and Rag1-Cre models to track the contribution of fetal progenitors to tissue resident macrophage (trMac) populations (Boiers et al., 2013; Epelman et al., 2014; Gomez Perdiguero et al., 2015; Hashimoto et al., 2013; Hoeffel et al., 2015), the contribution of IL7R-marked progenitors to the same populations has not been previously examined. To decipher the contribution of specific transient hematopoietic progenitors to adult trMacs, we compared trMac labeling across three lineage tracing models: Flk2-Cre, IL7R-Cre, and Rag1-Cre. We crossed mice expressing Rag1-Cre (Welner et al., 2009) or IL7R-Cre (Schlenner et al., 2010) to mTmG mice expressing a dual color fluorescent reporter (Muzumdar et al., 2007), thereby creating “Rag1Switch” and IL7RSwitch” models (Fig.1A) analogous to the previously described FlkSwitch mouse (Beaudin et al., 2016; Boyer et al., 2012; Boyer et al., 2011). In all models, all cells express Tomato (Tom) until Cre-mediated recombination results in the irreversible switch to GFP expression by that cell and all of its progeny (Fig.1B).

**Figure 1.**
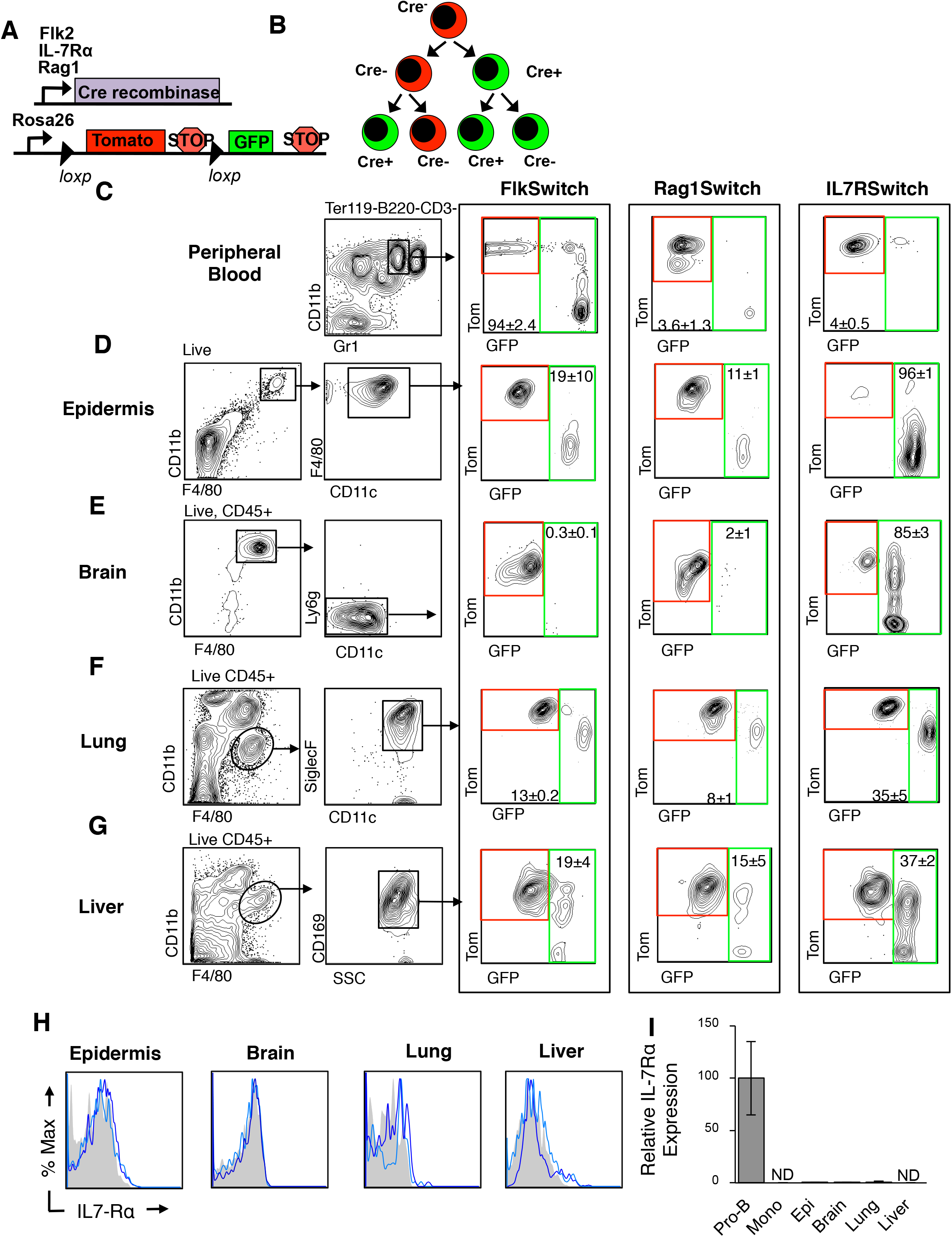
IL7R-Cre specifically labels adult tissue-resident macrophage populations. Representative flow cytometric analysis of reporter expression across different monocyte and macrophage populations in adult mice. Tom and GFP expression is highlighted by red and green boxes, respectively, in FlkSwitch, Rag1Switch, and IL7Rswitch models. Values indicate mean frequencies ± SEM of gated IL7R-Cre marked GFP+ populations. Plots and values are representative of 4-5 mice each representing three independent experiments. **A,** Schematic of the “Switch” models. Cre recombinase expression was controlled by either Flk2, IL7Rα, or Rag1 regulatory elements, respectively. Cre-driver mice were crossed to mice expressing a dual color reporter expressing either Tomato (Tom) or GFP, under control of the Rosa26 locus. Expression of Cre results in an irreversible genetic deletion event that causes a switch in reporter expression from Tomato to GFP. **B,** Schematic of Cre-mediated reporter switching in the “switch” models. All cells initially express Tomato. Expression of Cre results in an irreversible switch from Tomato to GFP expression. Once a cell expresses GFP, it can only give rise to GFP-expressing progeny. **C,** Representative flow cytometric analysis of reporter expression in circulating CD11b^hi^Gr^mid^ monocytes in the peripheral blood (PB) of adult FlkSwitch, Rag1Switch, and IL7RSwitch mice. **D,** Representative flow cytometric analysis of reporter expression in Langerhans cells (F4/80+CD11b+CD11c^mid^) in the epidermis of adult FlkSwitch, Rag1Switch, and IL7RSwitch mice. **E,** Representative flow cytometric analysis of reporter expression in microglia (CD45+F4/80^hi^CD11b^hi^Ly6g^−^CD11c^−^) in the brains of FlkSwitch, Rag1Switch, and IL7RSwitch adult mice. **F,** Representative flow cytometric analysis of reporter expression in lung alveolar macrophages (CD45^+^F4/80^hi^CD11b^mid^ SiglecF^+^CD11c^+^) of FlkSwitch, Rag1Switch, and IL7RSwitch adult mice. **G,** Representative flow cytometric analysis of reporter expression in Liver Kupffer cells (CD45^+^F4/80^hi^CD169^+^) in adult FlkSwitch, Rag1Switch, and IL7RSwitch mice. **H,** Lack of IL-7Rα surface expression in the Langerhans cells of the epidermis, brain microglia, lung alveolar macrophages, and liver Kupffer cells. For each tissue, IL-7Rα surface expression of gated population is shown for two representative mice in blue. Gray shaded area represents fluorescence-minus-one (FMO) control. **I,** Quantitative RT-PCR analysis of IL-7Rα expression in sorted bone marrow (BM)-derived B220+CD43+ ProB-cells and CD11b+Gr1+monocytes, and Langerhans cells of the epidermis, brain microglia, lung alveolar macrophages, and Liver Kupffer cells isolated from WT adult mice. Data represent mean ± SEM for 3 independent experiments. Values are normalized to Pro-B cells, set to 100. ND, not detected. Additional analyses of adult tissue myeloid populations can be found in Figure S1.

We compared reporter expression in fetal-derived trMacs of all three models to reporter expression in circulating peripheral blood (PB) monocytes (CD11b^hi^Gr1^lo^), the precursors of adult HSC-derived macrophages. As expected, Cre-driven GFP labeling of monocytes in IL7RαSwitch and Rag1Switch mice was less than 5% (Fig.1C) and paralleled the nominal labeling observed in adult HSCs and myeloid progenitors (Fig. S1A,B), as previously reported (Schlenner et al., 2010; Welner et al., 2009). Adult FlkSwitch mice exhibited high GFP labeling in PB monocytes (Figure 1C), as we previously reported (Boyer et al., 2011). Intriguingly, examination of Cre-driven reporter switching in adult trMacs revealed distinct GFP labeling patterns across all three lineage tracing models and across tissues. Within Langerhans cells (LCs) of the epidermis, around 20% of cells were labeled by GFP in the FlkSwitch model (Fig. 1D), consistent with previous reports (Gomez Perdiguero et al., 2015; Hoeffel et al., 2015). Low GFP labeling was observed in LCs of Rag1Switch mice (Fig. 1D), as described previously for skin and other tissue macrophages at E14.5 (Boiers et al., 2013). In sharp contrast, up to 96% of LCs expressed GFP in IL7RSwitch mice (Fig. 1D). A similar pattern was observed for microglia; consistent with a pre-HSC origin, virtually no reporter switching was observed in adult microglia of FlkSwitch mice (Fig.1E). Minimal microglia labeling was also observed in Rag1Switch mice. Remarkably, however, GFP-labeling of microglia was over 85% in IL7RSwitch mice (Fig.1E).

Examination of reporter expression in trMacs of the lung (alveolar macrophages, AMs) and the liver (Kupffer cells, KCs) yielded another labeling pattern across lineage tracing models, as compared to the LC and microglia (Fig.1F,G). In FlkSwitch and Rag1Switch mice, GFP labeling in AMs and KCs was low and roughly comparable to LCs (~8-19% GFP+ cells; Fig. 1D,F,G). Surprisingly, in IL7RSwitch mice, GFP labeling in AM and KC populations was substantially higher, ~40%, as compared to both FlkSwitch and Rag1Switch models (Fig.1F,G). Although labeling in these two populations was still considerably less than that of LC or microglia in the same mice (~85-96%), it was substantially higher than labeling of adult BM-derived myeloid populations (F4/80^lo^CD11b+) within the same tissues (<5%) (Fig. S1C-E). Comparison of Cre-mediated labeling between circulating BM-derived monocytes (4%) and trMacs (40-90%) in adult IL7RSwitch mice also revealed stark differences in labeling, supporting previous reports that adult BM-derived monocytes did not substantially contribute (at steady-state) to the trMac populations investigated, including microglia, Langerhans cells, Kupffer cells, and alveolar macrophages (Gomez Perdiguero et al., 2015; Hashimoto et al., 2013; Hoeffel et al., 2012; Sheng et al., 2015). Despite substantial IL7R-Cre-mediated labeling, IL-7Rα surface expression was undetectable in any adult trMacs surveyed (Fig. 1H). IL-7Rα message was also virtually undetectable in trMacs as compared to BM Pro-B-cells (Figure 1I). The robust labeling of adult trMacs in the IL7RαSwitch model therefore suggested IL-7Rα expression during an earlier window of macrophage development.

### YS hematopoiesis does not depend on IL7R

Yolk sac (YS) erythromyeloid progenitors (EMPs) have been proposed as a cell of origin for fetal trMac development, either through direct differentiation or through the myb-dependent generation of intermediate progenitors that seed the fetal liver (Gomez Perdiguero et al., 2015; Hoeffel et al., 2015; Hoeffel et al., 2012; Mass et al., 2016; Schulz et al., 2012). To determine whether IL7R-Cre labeling of adult trMac populations initiated in YS EMPs during embryonic development, we examined IL7R-Cre driven GFP labeling, IL-7Rα surface expression and IL-7Rα message in YS EMPs and YS macrophages. YS EMPs (Ter119-cKit+CD41+) at E9.5 were minimally labeled by IL7R-Cre, consistent with a complete absence of IL-7Rα surface expression (Fig 2A). IL-7Rα message was detectable at very high Ct values (Fig 2C, C’), and consequently we observed slightly more labeling in YS EMPs by E10.5 (Fig. 2A). However, increased reporter expression was not accompanied by surface expression (Fig. 2A’). Labeling of YS EMPs was comparable to background labeling observed in adult hematopoiesis (Schlenner et al., 2010). Few YS myeloid cells, defined by either CD11b or F4/80 expression within the whole CD45+ fraction, exhibited IL7R-Cre driven reporter expression (Fig. 2B) and they did not express surface IL-7Rα (Fig. 2B’). To determine whether IL-7R expression regulated YS hematopoiesis, we examined the frequency of YS EMPs and YS macrophages in IL7R−/− embryos. Neither YS EMPs nor YS macrophage frequency was significantly different between WT and IL7R−/− mice (Fig. 2D,E). Together, these data indicate that IL-7R is not required for the generation of YS progenitors or YS macrophages, and suggest that labeling of F4/80^hi^ trMacs by IL7R-Cre occurs later in trMac ontogeny.

**Figure 2.**
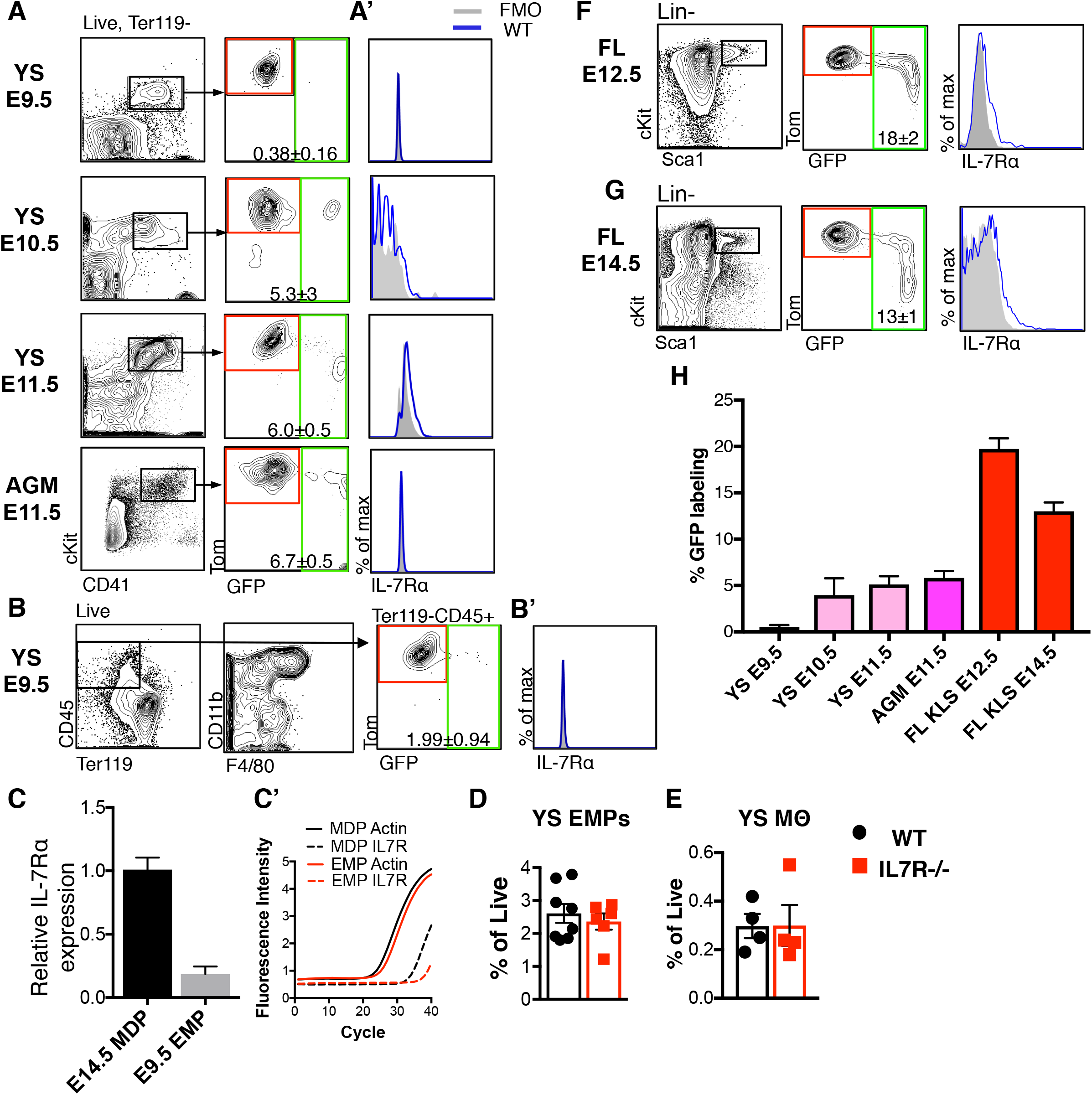
IL-7Rα is dispensable for YS hematopoiesis. **A, A’,** Representative FACS plots showing gating strategy, IL7R-Cre-driven GFP labeling, and IL-7Rα surface expression in Ter119-cKit+CD41+ erythromyeloid progenitors (EMPs) in the yolk sac (YS) at E9.5, E10.5, E11.5 and in the aorta-gonad-mesonephros region (AGM) at E11.5. Red and green boxes denote gates for Tom+ and GFP+ progenitors, respectively. Gray shaded areas in histograms indicate FMO control. Plots and values indicate mean frequencies ± SEM of gated IL7R-Cre marked GFP+ populations for 4-7 mice from at least two independent experiments. **B,** Representative FACS plots showing gating, IL7R-Cre-driven GFP labeling, and IL7Rα surface expression in CD45+Ter119-cells in the YS at E9.5. Adjacent plot shows CD11b and F4/80 expression within the CD45+Ter119-compartment. Red and green boxes denote gates for Tom and GFP expression of CD45+Ter119-cells, respectively. **C, C’,** Quantitative RT-PCR analysis of IL-7Rα mRNA in YS EMPs from E9.5. **(C)** ΔΔC^T^ values normalized to beta-Actin as compared to IL-7Rα expression levels in MDPs (CD115+cKit+Flk2+Ly6c-) isolated from E14.5 FL. (**C’**) Representative amplification plot of beta-actin (solid) and IL-7Rα (dashed) for quantitative RT-PCR of E14.5 MDPs and EMPs as described in (C). Data show mean ± SEM from 3 independent experiments. **D, E,** Quantification of the frequency of EMPs (D) and F4/80+ macrophages (E) in YS of IL7R^−/−^ and WT embryos at E9.5. Data show mean frequency ± SEM representing 3 independent experiments. **F, G,** Representative FACS plots showing gating, IL7R-Cre-driven GFP labeling, and IL-7Rα surface expression in cKit+Lin-Sca1+ stem and progenitor cells at E12.5 (F) and E14.5 (G). **H,** Quantification across development of the percent of IL7R-Cre driven reporter labeling (GFP+) in the phenotypically defined stem and progenitor populations described in A-G. Additional analysis of IL7R-Cre driven reporter labeling in fetal hematopoietic populations can be found in Figure S2.

### IL7Rα-Cre labels non-HSC progenitors in the fetal liver

As early YS progenitors and macrophages were minimally labeled by IL7R-Cre, we investigated labeling in putative progenitors slightly later in development. Labeling remained comparably low in both YS and AGM cKit+CD41+ EMPs at E11.5, and IL-7R surface expression was still undetectable in the EMPs within the FL (Fig. 2A, Fig. S2A). Higher IL7R-Cre driven labeling was observed in FL Lin-cKit+Sca1+ (KLS) progenitors by E12.5, but surface expression was still low (Fig. 2F, H). Labeling subsequently declined in E14.5 progenitors (Fig. 2G, H; p < 0.001), and all labeling in E14.5 progenitors was confined to the CD150-multipotent progenitor compartment (Fig. S2B). Consistent with labeling of upstream multipotent cells in the FL, comparable IL7R-Cre labeling was observed in all downstream progenitor and mature compartments (Fig S2C-G), with the exception of significant labeling in committed lymphoid progenitors (Fig. S2D) and mature lymphoid cells (Fig S2E). Labeled progenitors at E14.5 did not possess long-term multilineage reconstitution, confirming that they were not definitive HSCs (Fig. S2H). These data suggest that IL-7R reporter expression in late (E10.5) YS and AGM partially labels a transient progenitor that contributes to FL hematopoiesis but is not a definitive HSC.

### IL7R-Cre labeling of myeloid cells is tissue- and developmental stage-specific

Given that IL7R-Cre labeling in adult trMacs (Fig. 1) was considerably higher as compared to any progenitor we profiled across fetal development (Fig. 2H; Fig. S2), we reasoned that progenitor labeling alone could not entirely explain labeling observed in those compartments. To resolve this discrepancy, we next evaluated IL7R-Cre-driven labeling and IL-7Rα expression in myeloid-restricted precursors within peripheral tissues during FL stage development. We initiated our investigation at E12.5, the stage at which significant progenitor labeling by IL7R-Cre was initially observed (Fig. 2F,H). At this stage of development, macrophages derived from the extra-embryonic YS (F4/80^hi^CD11b^lo^) are already present in the tissues, but may be replaced by incoming FL-derived macrophage precursors or monocytes (F4/80^lo^CD11b^hi^, Fig. 3A). F4/80^lo^CD11b^hi^ macrophage precursors in fetal peripheral tissues are heterogeneous or low for Ly6c expression and express the fetal macrophage marker CD64 (Fig. S3A; (Hoeffel et al., 2015), suggesting their propensity to differentiate into macrophages.

**Figure 3.**
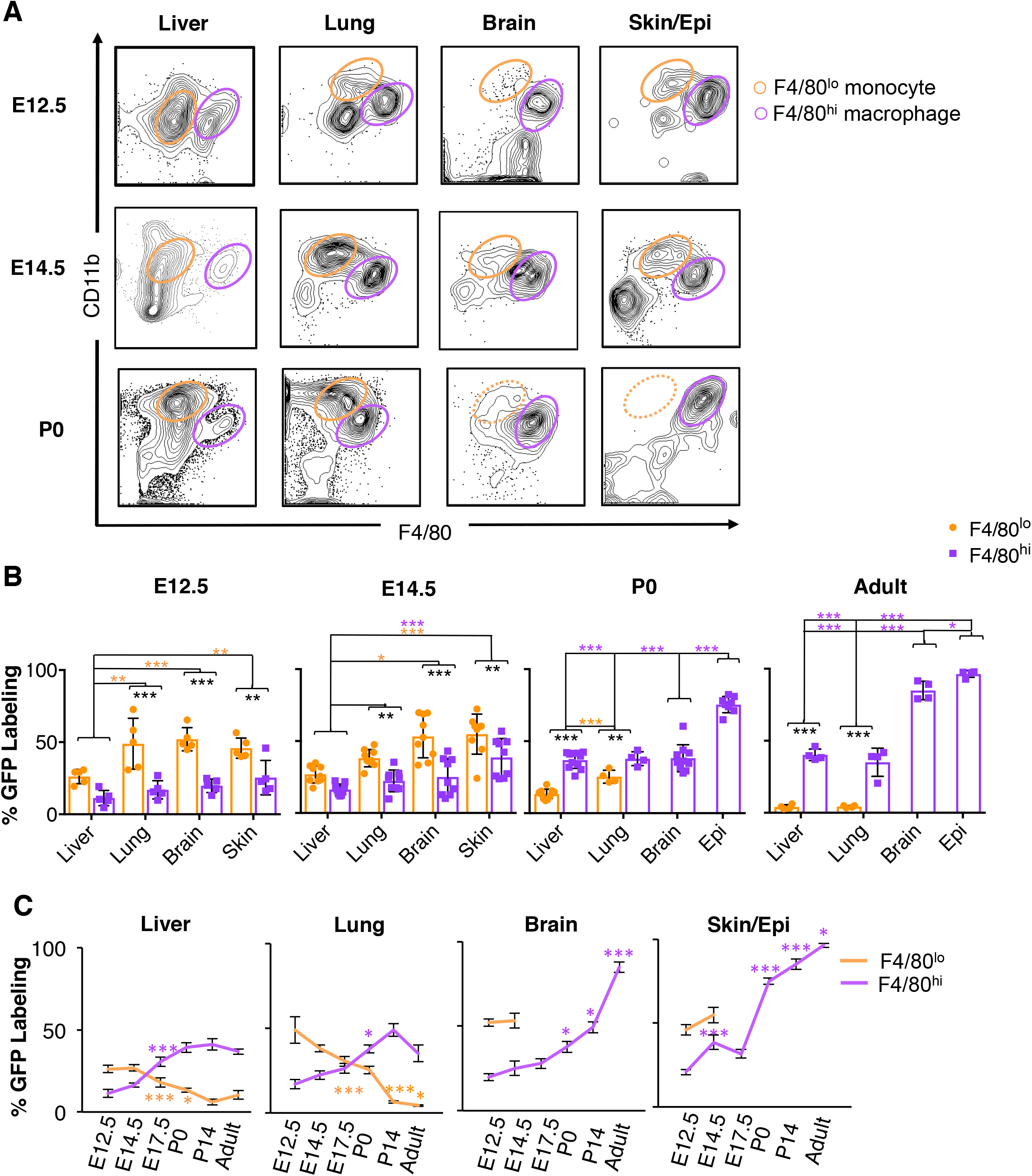
IL7R-Cre dynamically labels myeloid cells during tissue resident macrophage development. **A,** Representative flow cytometric analysis indicating gating for fetal liver (FL)-derived F4/80^lo^ monocytes (CD45+F4/80^lo^CD11b^hi^; orange gates) and previously seeded F4/80^hi^ macrophages (CD45+F4/80^hi^CD11b^mid^; purple gates) in different tissues at E12.5, E14.5 and P0. **B,** Quantification of IL7R-Cre driven GFP labeling within gated population indicated in (A) across different tissues at E12.5, E14.5, P0 and Adult (8-12 weeks). Orange asterisks denote cross-tissue differences in GFP labeling between F4/80^lo^ monocytes and purple asterisks denote cross-tissue differences in GFP labeling between F4/80^hi^ macrophages. Black asterisks denote differences in GFP labeling between F4/80^lo^ monocytes and F4/80^hi^ macrophages within each tissue. N = 4-11 representing at least 3 independent experiments. * P < 0.05, ** P < 0.005, *** P < 0.0005. **C,** Time course of IL7R-Cre driven GFP labeling in F4/80^hi^ macrophages and F4/80^lo^ monocytes shown in (B) across ontogeny. Asterisks indicate statistically significant differences between the percentage of GFP labeling at the previously measured timepoint, with purple asterisks denoting differences between the indicated F4/80^hi^ macrophages and orange asterisks denoting differences between F4/80^lo^ monocyte populations. Additional analysis of fetal myeloid populations can be found in Figure S3.

Analysis of IL7R-Cre driven reporter expression revealed two very clear patterns: first, GFP labeling at E12.5 was significantly higher in all F4/80^lo^ monocytes as compared to F4/80^hi^ macrophages across all tissues (Fig/ 3B, left plot/E12.5 data). Second, cross-tissue comparison revealed higher GFP labeling in F4/80^lo^ monocytes in brain, lung, and epidermis as compared to liver (Figure 3B, left plot/E12.5 data). Across all F4/80^hi^ macrophages, GFP labeling was highest in the skin (25%; P< 0.05 compared to liver F4/80^hi^ macrophages). A very comparable pattern emerged at E14.5; however, between E12.5 and E14.5, GFP labeling increased slightly across all F4/80^hi^ macrophage populations, but only significantly for the skin (Fig. 3C, Epidermis panel).

From late gestation into early neonatal development, the patterns of IL7R-Cre mediated GFP labeling among F4/80^hi^ macrophages and F4/80^lo^ monocytes were strikingly different, both between cell types and between tissues (Fig. 3B, C). Beyond E17.5, GFP labeling in liver and lung was higher in F4/80^hi^ macrophages as compared to F4/80^lo^ monocytes. This “switch” reflected a significant increase in GFP labeling of F4/80^hi^ macrophages between E14.5 and P0 and a concomitant decrease in F4/80^lo^ monocyte labeling (Fig. 3C), coincident with the HSC-dependent contribution to fetal monocytes later in gestation (Hoeffel et al., 2015; Yona et al., 2013). GFP labeling in lung and liver macrophages reached adult levels by P0 (Fig 3B, Figure 1F,G; P > 0.5). These data suggested that IL7R-Cre labeling was either occurring cell-autonomously in F4/80^hi^ macrophages or that IL7R-labeled fetal monocytes were gradually replacing tissue macrophage populations in the lung and liver during fetal hematopoiesis.

In contrast to lung and liver, progression of GFP labeling within the skin and brain across ontogeny relayed a different pattern. Monocytes were barely detectable in the epidermis and brain after E14.5 (Fig. 3A, bottom right panels), as previously described (Hoeffel et al., 2015). GFP labeling of skin macrophages increased robustly from E17.5 to P0 (Fig. 3C, skin), and continued to increase postnatally (Fig. 3B,C). Similarly, microglia labeling increased steadily across development, but had still not reached adult levels by postnatal day 14 (P14) (Fig. 3B,C). The disparate GFP labeling that we observed across different tissues suggested a differential involvement of IL-7R in the development of different trMac populations across ontogeny.

### IL-7Rα expression is dynamically regulated during fetal tissue macrophage development

To gain insight into the observed pattern of IL7R-Cre mediated GFP labeling within F4/80^lo^ monocyte and F4/80^hi^ macrophage populations across development, we probed IL-7Rα message levels beginning at E14.5 (Fig. 4A). Across tissues, F4/80^lo^ monocytes expressed low but consistently detectable levels of IL-7Rα message relative to fetal liver monocyte dendritic progenitors (MDPs) that are known to robustly express IL-7Rα message (Hoeffel et al., 2015) Fig. 4A). Circulating peripheral blood Ly6c+ monocytes also expressed detectable IL-7Rα message and were labeled by IL7R-Cre to a similar degree as compared to liver F4/80^lo^ monocytes (Fig S3C,D). Increased IL-7Rα mRNA in peripheral tissue F4/80^lo^ monocytes corresponded with higher Cre-mediated GFP expression (Fig. 4A), as GFP labeling increased in monocytes once they had exited the liver at E14.5 (Fig. 3B). Lung and skin monocytes with the highest GFP labeling also expressed the highest levels of IL7Rα mRNA. In comparison, and consistent with significantly lower Cre-mediated GFP labeling, IL-7Rα message was virtually undetectable in F4/80^hi^ macrophages (Figure 4A) during the prenatal period. Postnatally, at P14, IL-7Rα message levels continued to be negligible in macrophages of the lung and liver, but microglia and LCs expressed IL-7Rα message (Fig. 4B), driving continued postnatal Cre-mediated recombination in those tissues. These data confirmed the fidelity of the IL7R-Cre model and revealed IL7R expression by fetal myeloid cells during development.

**Figure 4.**
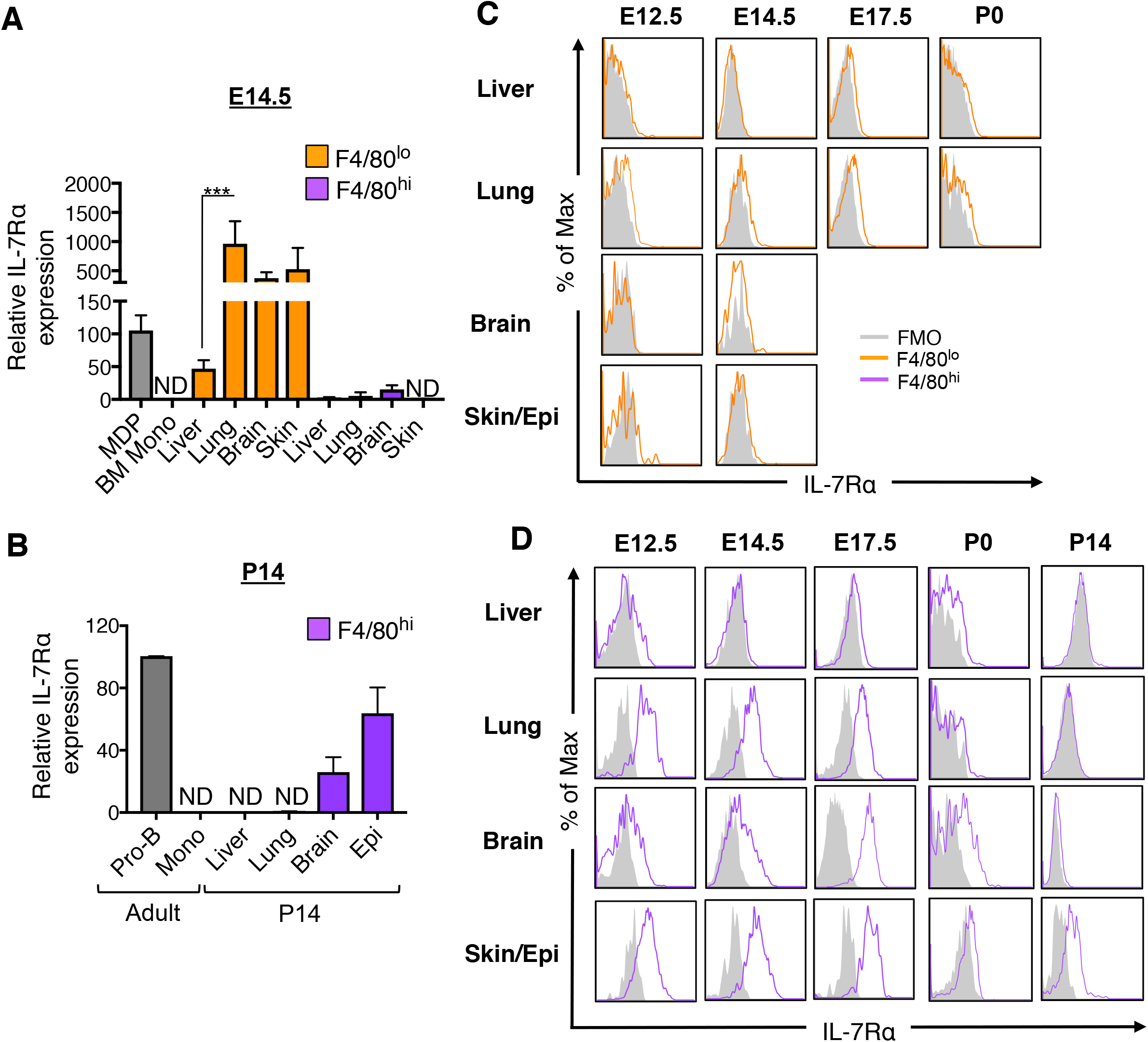
IL-7Rα message and surface expression are dynamically regulated during macrophage development. **A,** Quantitative RT-PCR analysis of IL-7Rα mRNA in monocytes (F4/80^lo^CD11b^hi^) and macrophages (F4/80^hi^CD11b^lo^) isolated from liver, lung, brain, and skin of E14.5 fetuses. Fetal liver macrophage dendritic cell precursors (MDP; CD115+cKit+Flk2+Ly6c-CD11b-) and adult BM monocytes (CD11b+Gr1mid) were used as positive and negative controls, respectively. Data shown are mean ± standard error of ΔΔCT values calculated for IL7Rα and Actin and normalized to MDPs (set to 100) across three independent experiments. **B,** Quantitative RT-PCR analysis of IL-7Rα mRNA in monocytes (F4/80^lo^CD11b^hi^) and macrophages (F4/80^hi^CD11b^lo^) isolated from liver, lung, brain, and epidermis of neonatal mice at P14. Adult BM ProB cells (B220+CD43+) and BM monocytes (CD11b+Gr1+) were used as positive and negative controls, respectively. Data shown are mean ∓ standard error of ΔΔCT values calculated for IL-7Rα and beta-actin and normalized to BM ProB cells (set to 100) across three independent experiments. ND, not detected **C, D** Representative flow cytometric analysis of IL-7Rα surface expression in F4/80^lo^ monocytes (CD45+F4/80^lo^CD11b^hi^; **C)** and F4/80^hi^ macrophages (CD45+F4/80^hi^CD11b^mid^; **D)** in different tissues at embryonic day (E)12.5, E14.5, E17.5, postnatal day (P)0 and P14. Gray shaded area indicates FMO control. Plots are representative of analysis in 5-6 mice from three independent experiments. Analysis of common gamma chain expression in fetal tissue macrophages can be found in Figure S4.

To ascertain the relationship between IL7R-Cre mediated GFP labeling and IL-7Rα surface expression in developing macrophages during fetal development, we profiled surface expression from E12.5-P14. As GFP labeling and IL7Ra mRNA expression was higher in F4/80^lo^ monocytes as compared to F4/80^hi^ macrophages during fetal development, we expected to observe monocytes displaying surface IL7R. Unexpectedly, F4/80^lo^ monocytes never displayed detectable IL-7Rα surface protein at any timepoints examined (Fig. 4C). Surprisingly, robust IL-7Rα surface protein was instead observed on prenatal F4/80^hi^ macrophages in peripheral tissues (lung, brain, and skin), whereas minimal surface expression was observed on F4/80^hi^ macrophages in the liver (Fig. 4D). IL-7Rα surface protein on peripheral F4/80^hi^ trMacs was observed beginning at E12.5, and peaked at E17.5 (Figure 4B, middle panel), despite significantly lower GFP labeling as compared to F4/80^lo^ monocytes (Fig. 3B,C). The vast majority (>90%) of IL-7Rα-expressing macrophages in the brain, lung, and skin also coexpressed the common γ chain (Fig. S4). By P0, macrophages in the liver and lung ceased to express IL-7Rα, concomitant with a plateau in GFP labeling by birth (Fig.3C). In contrast, some proportion of macrophages in the brain and epidermis continued to express IL-7Rα protein at the surface postnatally, albeit at lower levels, and IL-7Rα surface expression was detectable on epidermal macrophages as late as P14. Continued IL-7Rα expression in these particular tissues was consistent with continued postnatal labeling by IL7R-Cre (Fig.3C) and IL-7Rα mRNA expression (Fig. 4B). These data therefore revealed tightly regulated and highly coordinated expression of IL-7Rα in developing macrophages across different tissues.

### IL-7Rα -expressing monocytes give rise to IL-7R+ macrophages ex vivo

We hypothesized that F4/80^lo^CD11b^hi^ monocytes upregulate IL-7Rα message as they exit the liver and migrate to and enter fetal tissues, where they differentiate into F4/80^hi^ macrophages. IL-7Rα surface expression is then rapidly switched on as F4/80^lo^ monocytes differentiate into F4/80^hi^ macrophages. To test our hypothesis, we isolated F4/80^lo^ monocytes from the fetal liver at E14.5 and differentiated them into F4/80^hi^ macrophages *ex vivo* in the presence of M-CSF. F4/80^lo^CD11b^hi^cells were sorted from FL based on Ly6C expression (Fig. 5A) and cultured with 20ng M-CSF/mL for 5 days. FL monocytes expressed detectable IL-7Rα message (Fig. 4A), but IL-7R surface expression was not observed (Fig 5C, C’). M-CSF induced the differentiation of Ly6c^hi^CD11b^hi^ cells to F4/80^hi^CD11b+ macrophages that expressed CD64 after only 1 day in culture, and macrophage differentiation plateaued at 3 days (Fig 5C,E, E’). Remarkably, differentiated F4/80^hi^ cells also upregulated IL-7Rα expression on the cell surface (Fig. 5C, C’). IL-7Rα was co-expressed with the common gamma chain, indicating expression of a functional IL-7R receptor (Fig. 5D, D’). In contrast to Ly6c^hi^ cells, Ly6c^lo^CD11b^hi^ FL monocytes displayed limited differentiation into F4/80^hi^ macrophages, even after 5 days in culture with M-CSF (Fig. S3F), and both MDP and Ly6c^lo^ monocytes displayed significantly higher Cre-mediated labeling as compared to Lyc6^hi^ monocytes (Fig. S3C), suggesting distinct developmental pathways.

**Figure 5.**
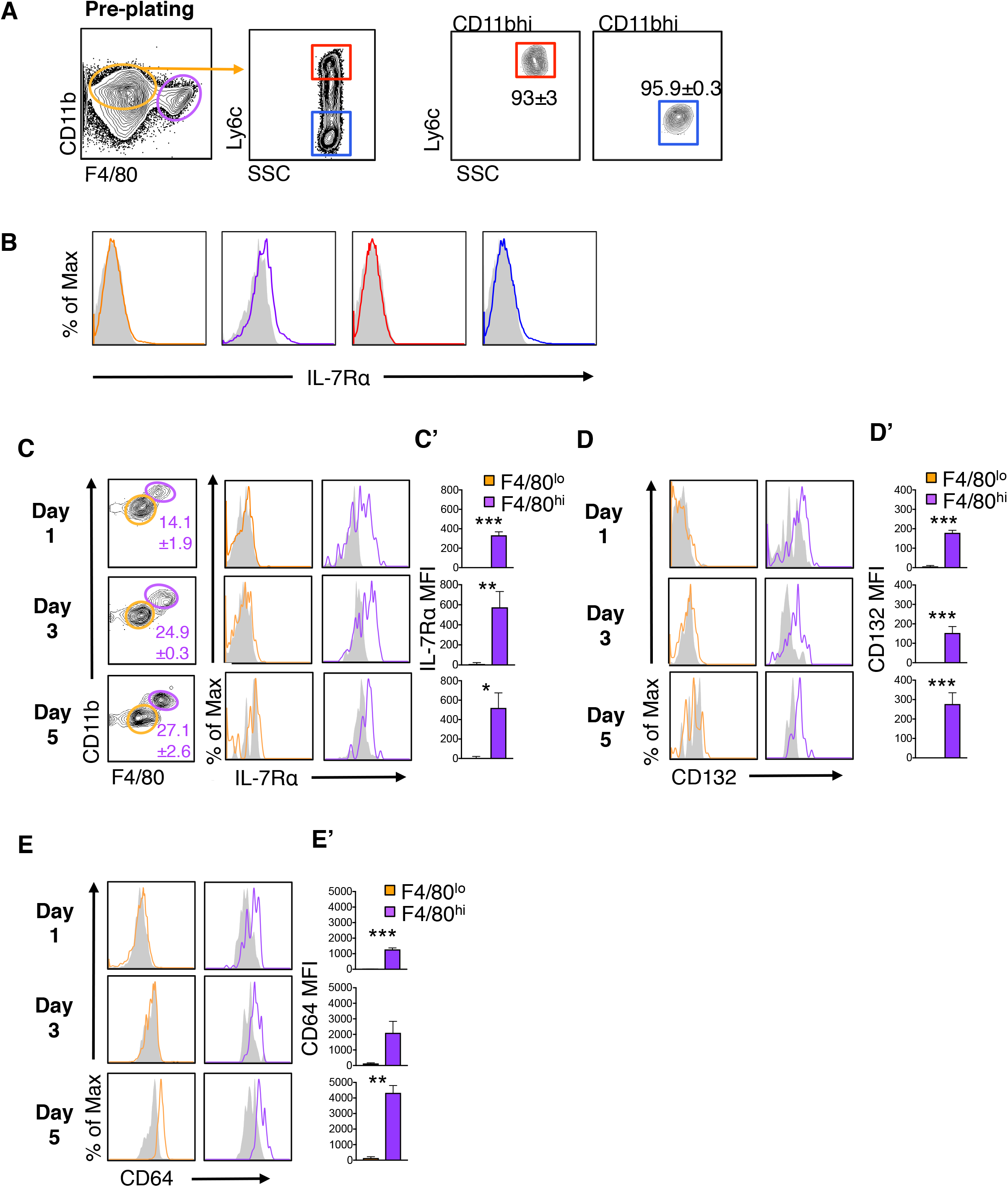
Fetal liver monocytes differentiate into IL-7R-expressing macrophages ex vivo. **A,** Representative gating of F4/80^lo^CD11b^hi^monocytes (orange gates) from CD11b-enriched fetal liver (FL) at E14.5, gated on Ly6c expression (Ly6c^hi^, Red; Ly6c^lo^, Blue). Adjacent plots represent post-sort purity analysis of FL CD11b^hi^F4/80^lo^ Ly6c^hi^ and Ly6c^lo^ monocytes with mean ± SEM frequency of sorted populations in triplicate from 3 independent experiments. **B,** Representative FACS plots show surface IL-7Rα expression on macrophage (F4/80^hi^) and monocyte (F4/80^lo^CD11b^hi^) populations as a function of Ly6c expression, color-coded as gated in (A). **C-E,** Representative FACS plots showing gating of monocyte (F4/80^lo^CD11b+) and macrophage (F4/80^hi^CD11b+) populations and expression of IL-7Rα **(C)**, CD132 **(D)**, and CD64 **(E)** on gated populations following 1, 3, or 5 days of culture of sorted F4/80^lo^CD11b^hi^Ly6c^hi^ monocytes with 20 ng m-CSF. Mean fluorescence intensity (MFI; **C’, D’, E’**) was calculated as the geometric mean minus the FMO for each sample. Gray shaded area indicates FMO control. Values indicate mean frequencies ± SEM of gated populations for 6-8 different samples in triplicate from 3 independent experiments. *. P < 0.05, ** P< 0.01, *** P < 0.001, Student’s T-test.

These data reveal that Ly6c^hi^CD11b^hi^ cells in the fetal liver that express IL-7Rα message (Fig. 4A; Fig S3D) but not surface IL-7R surface protein (Fig. S3E) have the capacity to differentiate into F4/80^hi^ macrophages, and that this differentiation is accompanied by upregulation of surface IL-7R expression.

### IL-7Rα regulates tissue resident macrophage development

The robust and dynamic expression of IL-7R by developing macrophages themselves suggested that IL-7R regulates fetal trMac development from precursors within fetal tissues. To test this hypothesis, we injected the highly specific IL-7Rα monoclonal blocking antibody, A7R34, or an IgG2A control, into pregnant mice during fetal development (Fig. 6A). Injection of A7R34 during pregnancy completely blocks IL-7R signaling, as evidenced by deletion of Peyer’s patches in developing embryos following a single injection (Hashizume et al., 2008; Yoshida et al., 1999). We injected pregnant WT mice with A7R34 or IgG2A control at E13.5 and E15.5, coincident with robust IL-7Rα expression on fetal macrophages (Fig. 4B), and examined cellularity of macrophages in the epidermis, lung, liver, and brain in neonates 9 days later (Fig. 6A). Macrophage cellularity in the epidermis, liver, and lung was significantly reduced following this temporally-limited blockade of IL-7Rα signaling, whereas microglia were unchanged (Fig. 6B). Coincidentally, limited IL-7R blockade resulted in a trend towards increased CD11b^hi^F4/80^lo^ monocytes in the fetal liver (p = 0.07) as well as a significant reduction in F4/80^lo^ monocytes in the lung (p < 0.05). These data provide additional evidence that IL-7R is regulating the transition from F4/80^lo^ macrophage precursor to F4/80^hi^ macrophage in peripheral tissues.

**Figure 6.**
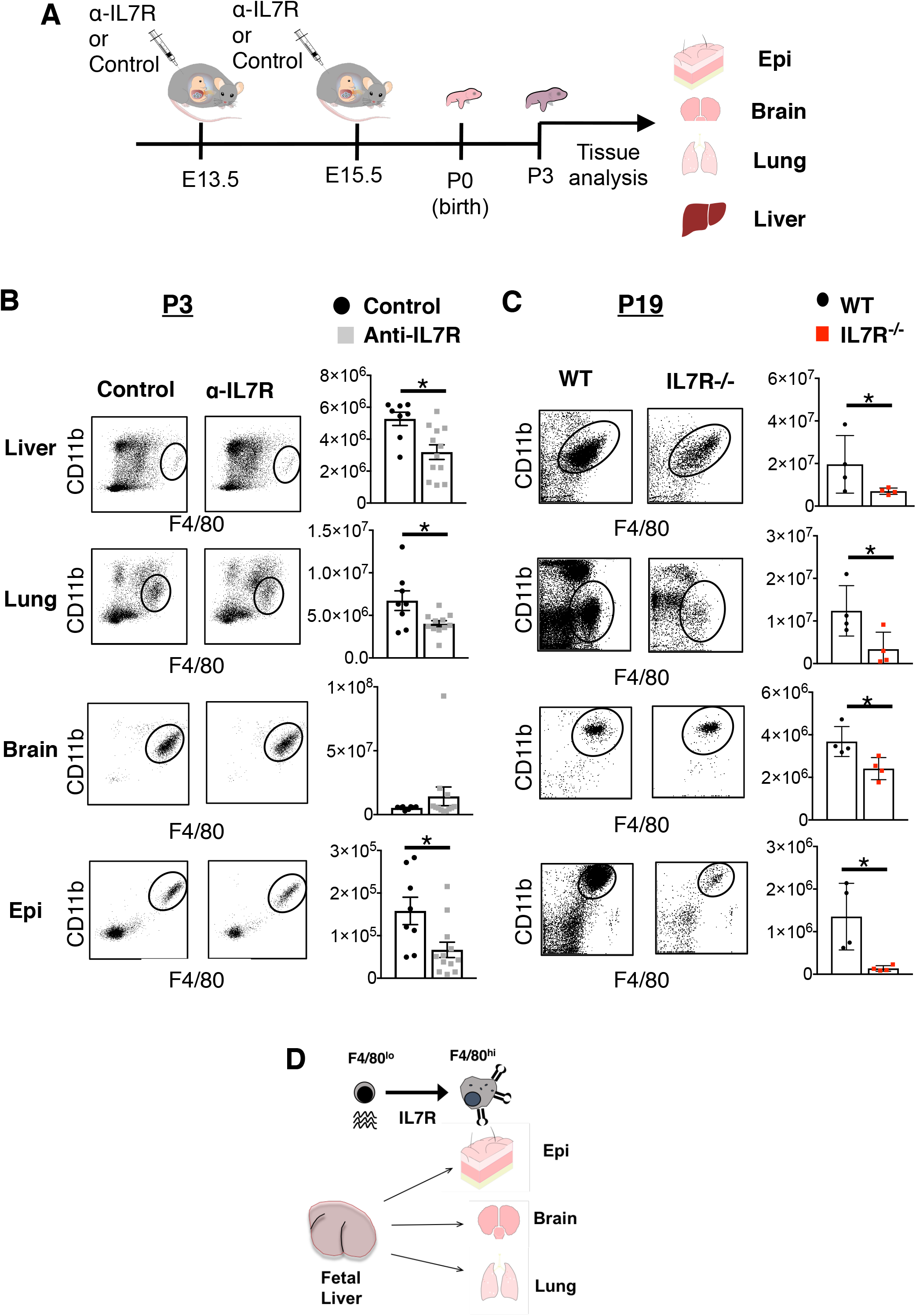
IL-7R regulates tissue resident macrophage development. **A,** Schematic illustrating the timeline for injection and analysis of the IL7R blocking antibody, A7R43, and IgG2A control. Timed-mated WT mice were injected at embryonic day (E)13.5 and E15.5 with 600 mg of the IL-7Rα blocking antibody or equivalent control injection. Pups were analyzed 9 days after final injection and cellularity of macrophages in the liver, lung, brain, and epidermis were determined. **B,** Representative FACS plots and associated quantification of cellularity of tissue macrophages (F4/80^hi^CD11b^lo^) in the liver, lung, brain, and epidermis of neonates examined nine days following maternal injections of the A7R43 IL-7Rα blocking antibody or IgG2A control administered at embryonic day (E)13.5 and E15.5. N = 8-11 representing three independent experiments. *, P< 0.05. **C,** Representative FACS plots and associated quantification of cellularity of tissue macrophages (F4/80^hi^CD11b^lo^) in the liver, lung, brain, and epidermis of IL7R^−/−^ and WT neonates at postnatal day (P) 19. N = 4 representing four independent experiments. *, P< 0.05. **D,** A model for dynamic IL-7Rα expression during fetal macrophage development. Fetal liver F4/80^lo^ monocytes express IL-7Rα message, and mRNA expression is upregulated as F4/80^lo^ monocytes exit the liver and transit to peripheral tissues. However, upregulation of message is not associated with measurable surface expression in F4/80^lo^ monocytes. IL7Rα surface expression only occurs upon differentiation of monocytes into F4/80^hi^ macrophages in the tissues, at which point message levels dissipate. Continued message and surface expression postnatally within microglia and Langerhans cells account for higher levels of GFP labeling in those tissues, whereas GFP labeling is equilibrated by birth in the alveolar macrophages of the lung and Kupffer cells of the liver.

We further investigated trMac cellularity in IL7R^−/−^ mice, which have a germline deletion of IL-7Rα (Peschon et al., 1994). Although IL-7R deletion did not affect YS hematopoiesis (Fig. 2) examination at P19 revealed that IL-7R deletion drastically reduced cellularity of trMacs in all tissues (Fig. 6C). The most severe reductions were observed in the lung and epidermis (Fig. 6C), consistent with macrophages in those tissues expressing the highest and most sustained levels of IL-7R across development. Together with the IL-7R blocking experiments, these data indicate that IL-7R plays a unique role in the establishment of trMacs during perinatal development.

## Discussion

Our investigation has revealed a novel role for IL-7R in fetal myeloid development, and specifically in the generation of fetal-specified trMacs. Tracing the progeny of cells mapped by the lymphoid marker IL-7Rα revealed that trMacs derived from fetal hematopoiesis were distinctly marked by IL-7Rα expression history (Fig. 1). In adult BM hematopoiesis, IL-7Rα expression exclusively marks lymphoid cells (Schlenner et al., 2010), and the many functions of IL-7R signaling in lymphopoiesis are well-delineated (Fry and Mackall, 2005). We confirmed that adult trMacs do not express IL-7Rα message or protein (Fig. 1H,I), and that both YS EMPs and adult BM-derived myeloid cells are minimally labeled by IL7R-Cre (Fig. 1C, 2A, S1). Instead, our developmental analysis revealed that labeling of trMacs by IL7R-Cre occurred as a result of dynamic stage- and tissue-specific expression of IL-7R during macrophage development within fetal tissues (Figs. 3,4). We showed that the CD11b^hi^Ly6c^hi^ fetal liver monocytes rapidly upregulated surface IL-7R as they differentiated into F4/80^hi^ macrophages ex vivo (Fig. 5). Using two different loss-of-function approaches, we further demonstrated that blocking IL-7R signaling during the window of robust IL7R expression by developing trMacs impairs their establishment (Fig. 6). By revealing a role for IL7R in trMac development, our data define a new signaling mechanism regulating trMac establishment in late gestation and identify a function of IL7R during fetal development that extends beyond regulation of the lymphoid lineage.

The comparison of trMac labeling across three different lineage tracing models based on lymphoid markers associated with both increasingly restricted lymphoid potential (Flk2, Rag1, IL-7Rα; Fig. 1) (Adolfsson et al., 2005; Forsberg et al., 2006; Igarashi et al., 2002; Kondo et al., 1997) as well as overlapping expression in FL progenitors (Beaudin et al., 2016; Boiers et al., 2013) allowed us to exclude the possibility that labeling of adult trMacs by IL7R-Cre reflected inheritance from an earlier lymphomyeloid progenitor. A clear example is epidermal langerhans cells: if labeling of LC was due to derivation from a lymphomyeloid progenitors, LC would be robustly labeled not only by IL7R-Cre, but also by Flk2-Cre and Rag1-Cre, as those genes are expressed in IL7R+ FL progenitors (Boiers et al., 2013). The high degree of IL7R-Cre labeling of trMacs was similarly not reflected in YS EMPs or YS macrophages (Fig. 2, Fig S2) and embryonic YS EMPs did not express significant IL-7Rα message nor IL-7R surface protein. Instead, our tracking of IL7R-Cre mediated labeling (Fig. 3), IL-7Rα mRNA levels (Fig. 4A) and IL-7Rα surface expression (Fig. 4C), revealed dynamic regulation of IL-7Rα expression by developing macrophage precursors and macrophages themselves during later stages of fetal hematopoiesis. Furthermore, blocking IL-7R during this limited window of development (E13.5-E15.5), when IL-7R was robustly expressed by fetal macrophages (Fig. 4D), impaired establishment of those populations (Fig. 6B). In contrast, IL-7R deletion had no effect on YS EMP cellularity or the development of YS macrophages during embryonic hematopoiesis (Fig. 2D, E), but resulted in significant impairments later in trMac development (Fig. 6C). Together, these data suggest that IL7R expression is important for the development of macrophages within tissues later in gestation, possibly during later waves of macrophage seeding (De et al., 2018; Ferrero et al., 2018; Tan and Krasnow, 2016).

Fetal F4/80^hi^ macrophages expressed negligible IL-7Rα message levels (Fig. 4A) yet expressed robust surface IL7R protein (Fig. 4D) and acquired IL7R-Cre labeling across fetal development (Fig. 3C). In lung and liver, for example, increased GFP labeling in F4/80^hi^ trMacs across fetal development did not reflect Cre-driven reporter expression, as trMacs did not express IL-7Rα message during fetal development (Fig. 4A). The most parsimonious explanation for increased labeling is the initiation of Cre recombination in IL7R-marked F4/80^lo^ macrophage precursors with subsequent inheritance of the floxed allele by F4/80^hi^ trMacs upon differentiation. F4/80^hi^ trMac differentiation is then coincident with the rapid translation of IL-7Rα message into IL-7Rα protein and surface display (Fig. 6D). Indeed, our ex vivo differentiation assay revealed that Ly6c+ FL monocytes expressing IL-7Rα mRNA (Fig. 4C; Fig. S3D) differentiate into F4/80^hi^ macrophages that upregulate surface IL-7R protein (Fig. 5). By tracking GFP expression within fetal myeloid cells in the IL7R-Cre model, we observed the contribution or replacement of GFP-labeled F4/80^lo^ precursors to F4/80^hi^ trMacs in the lung and liver, as evidenced by increased labeling in F4/80^hi^ macrophages across ontogeny (Fig. 3C). Whereas trMacs of the lung and liver exhibited robust labeling (~40%) that had plateaued at birth (Fig. 3A-C), IL-7Rα mRNA expression (Fig. 4B, D; (Mass et al., 2016)) and Cre-mediated labeling of LC and microglia continued postnatally, leading to almost complete labeling by IL7R-Cre in adulthood. In vivo, the rapid and dynamic regulation of IL-7R protein surface expression may reflect a response to ligand in tissues, as surface expression of IL-7R is regulated by IL-7 availability (Clark et al., 2014; Wei et al., 2000). The lower surface expression of IL-7R on fetal liver F4/80^hi^ macrophages as compared to peripheral tissues (Fig. 4D) in vivo may also reflect surface regulation by immediate sources of IL-7 ligand in the fetal liver. Together, our analysis provides additional support for the contribution of FL F4/80^lo^ macrophage precursors to specific trMac compartments during late gestation (Guilliams et al., 2013; Hoeffel et al., 2015; Rantakari et al., 2016; Tan and Krasnow, 2016), and reveal dynamic regulation of IL7R expression during macrophage differentiation.

Our analysis of tissue macrophage development in the IL7R KO mouse and in response to physiological blockade of IL-7R reveal IL-7R as a novel regulator of tissue macrophage development. Blocking IL-7Rα function during a narrow fetal window (E13.5 - E15.5) negatively affected cellularity of liver, lung, and epidermal macrophages, with no effect on microglia. In contrast, germline deletion of IL-7Rα in the IL7R^−/−^ mouse significantly impaired trMac cellularity across all tissues, including microglia, examined at postnatal day 19 (Fig. 6C). The different effect of total knockout and transient blockade of IL-7Rα on trMac development suggests that the window of dependency on IL7R signaling differs between different tissues. Furthermore, the effect of transient blockade of IL7R signaling at a stage when fetal macrophages themselves express robust surface levels of IL7R (e.g. late gestation) suggests that IL7R is directly regulating macrophage development in peripheral tissues. In contrast, deletion of IL7R had no effect on YS EMPs or YS macrophage generation, affirming that IL7R plays a more critical role in fetal macrophage generation at later stages of hematopoiesis. Regardless of whether IL7R+ fetal progenitors contribute to the trMac pool, the uniform and robust IL7R surface expression on fetal trMac cells themselves was highly surprising, and contributes to accumulating evidence that fetal hematopoietic lineage specification is considerably less constrained than adult hematopoiesis (Beaudin et al., 2016; Mebius et al., 2001; Notta et al., 2016).

## Methods

### Mice

All animals were housed and bred in the AALAC accredited vivaria at UC Santa Cruz or UC Merced and group housed in ventilated cages on a standard 12:12 light cycle. All procedures were approved by the UCSC or the UC Merced Institutional Animal Care and Use (IACUC) committees. IL7Rα-Cre (Schlenner et al., 2010), Rag1-Cre (Welner et al., 2009), and Flk2-Cre (Benz et al., 2008) mice, obtained under fully executed Material Transfer Agreements, were crossed to homozygous Rosa26^mTmG^ females (Muzumdar et al., 2007) to generate “switch” lines, all on the C57Bl/6 background. WT C56Bl/6 mice were used for controls and for all expression experiments. Adult male and female mice were used randomly and indiscriminately, with the exception of the FlkSwitch line, in which only males were used because many female mice do not carry a Cre allele. Similarly, mice for developmental analysis were used indiscriminately without knowledge of gender.

### Tissue and cell isolation

Mice were sacrificed by CO_2_ inhalation. Gravid uteri removed, and individual embryos dissected. Adult liver, lung, brain, and skin were isolated and treated with 1x PBS(+/+) with 2% serum, 0.2-1mg/ml collagenase IV (Gibco) with or without 100U/ml DNase1 for 20 minutes to 2 hours. For adult epidermis isolation, ears were first incubated with 1X PBS (+/+) containing 2.4 mg/ml dispase (Gibco) to separate the epidermis, followed by 2 hours incubation of the epidermis with 1/mg/mL collagenase. Following incubation, all tissues were passaged through a 16g needle or a 19g needle 10X, and then filtered through a 70 μM filter.

### Flow Cytometry

Cell labeling was performed on ice in 1X PBS with 5 mM EDTA and 2% serum. Antibodies used are listed in the supplementary data table 1. Analysis was performed on BD FACS Aria II at University of California-Santa Cruz, and BD FACS Aria III and the University of California-Merced and analyzed using FlowJo.

### Transplantation Assays

Transplantation assays were performed as previously described. (Beaudin et al., 2014; Beaudin et al., 2016; Smith-Berdan et al., 2015; Ugarte et al., 2015) Briefly, sorted Tom+ or GFP+ KLS cells were isolated from IL7RαSwitch or Rag1Switch E14.5 fetal liver donors. WT recipient mice aged 8-12 weeks were sublethally iradiated (750 rad, single dose). Under isofluorane-induced general anesthesia, sorted cells were transplanted IV. Recipient mice were bled 4, 8, 12, and 16 weeks post transplantation via tail vein and peripheral blood was analyzed for donor chimerism by means of fluorescence profiles and antibodies to lineage markers. Long-term multilineage reconstitution was defined as chimerism in both the lymphoid and myeloid lineages of > 0.1% at 16 weeks post-transplantation.

### Ex vivo macrophage culture

E14.5 fetal livers from WT embryos were harvested and homogenized via trituration. CD11b+ cells were enriched by positive selection with CD11b biotin-conjugated antibodies and streptavidin microbeads, using LS columns (Miltenyi). F4/80^lo^ CD11b^hi^ cells were sorted based on Ly6c expression and cultured in a 96-well plate (DMEM, 20% FBS, 1 mM Sodium Pyruvate, 10 mM HEPES, 0.1 mM 2-Mercaptoethanol, and 50 mg/mL primocin, Life technologies). Cells were cultured in triplicate in the presence of 20 ng/mL M-CSF and analyzed at three different time points (Days 1, 3, and 5) for the presence of F4/80^hi^ CD11b+ macrophages and for surface expression of IL7R, common gamma chain (CD132), and CD64.

RNA Isolation and Quantitative Real-time Polymerase Chain Reaction (qRT-PCR) Analysis: RNA isolation from sorted cells was accomplished using the RNeasy mini kit (Qiagen, Hilden, Germany), and cDNA was reverse transcribed from RNA (High capacity cDNA reverse transcription kit, ThermoFisher Scientific, Massachussetts). TaqMan probes (TaqMan Gene Expression Assay, ThermoFisher Scientific) were used for qRT-PCR analysis on a StepOnePlus Real-Time PCR System (ThermoFisher Scientific) in comparative CT mode. Samples were run in triplicate and were run with positive (CD43+ B220+ Pro B-cells), negative (Gr1+ CD11b+ BM monocytes), and no cDNA controls.

### Quantification and Statistical Analysis

Number of experiments, n, and what n represents can be found in the legend for each figure. Statistical significance was determined by two-tailed unpaired student’s T-test. All data are shown as mean ± standard error of the mean (SEM) representing at least three independent experiments.

## Acknowledgments

We thank Drs. Hans-Reimer Rodewald and Susan M. Schlenner for the IL7Rα^Cre^ strain; Drs. Patricia Ernst and Terri Rabbitt for the Rag1^Cre^ strain; Dr T. Boehm for the Flt3^Cre^ strain; Bari Nazario and the UCSC Institute for the Biology of Stem Cells for flow cytometry support; and David Gravano and the UC Merced Stem Cell Instrumentation Foundry for flow cytometry support.

## Funding

A.E.B. is the recipient of an NHLBI Mentored Career Development Award (K01HL130753). This work was supported by an NIH/NIDDK award (R01DK100917), an Alex’s Lemonade Stand Foundation Innovation Award, and an American Asthma Foundation Research Scholar award to ECF; by CIRM SCILL grant TB1-01195 to TM via San Jose State University; and by CIRM Facilities awards CL1-00506 and FA1-00617-1 to UCSC. ECF is the recipient of a CIRM New Faculty Award (RN1-00540) and an American Cancer Society Research Scholar Award (RSG-13-193-01-DDC).

## Author contributions

A.E.B. and E.C.F. conceived of the study, designed the experiments, and co-wrote the paper. A.E.B., T.M., G.L., A.W., and C.V. performed experiments and analyzed data. All authors reviewed the manuscript.

## Competing interests

The authors have no conflicting financial interests.

